# Targeting Corticotropin-Releasing Hormone Receptor Type 1 (CRHR1) Neurons: Validating the Specificity of a Novel Transgenic *Crhr1*-FlpO Mouse

**DOI:** 10.1101/2024.11.08.622735

**Authors:** Mason Hardy, Yuncai Chen, Tallie Z. Baram, Nicholas J. Justice

## Abstract

**Introduction:** Corticotropin-releasing hormone (CRH) signaling through its cognate receptors, CRHR1 and CRHR2, contributes to diverse stress-related functions in the mammalian brain. Whereas CRHR2 is predominantly expressed in choroid plexus and blood vessels, CRHR1 is abundantly expressed in neurons in discrete brain regions, including the neocortex, hippocampus and nucleus accumbens. Activation of CRHR1 influences motivated behaviors, emotional states, and learning and memory. However, it is unknown whether alterations in CRHR1 signaling contribute to aberrant motivated behaviors observed, for example, in stressful contexts. These questions require tools to manipulate CRHR1 selectively. Here we describe and validate a novel *Crhr1*-FlpO mouse.

**Methods:** Using bacterial artificial chromosome (BAC) transgenesis, we engineered a transgenic mouse that expresses FlpO recombinase in CRHR1-expressing cells. We used two independent methods to assess the specificity of FlpO to CRHR1-expressing cells. First, we injected *Crhr1*-FlpO mice with Flp-dependent viruses expressing fluorescent reporter molecules. Additionally, we crossed the *Crhr1*-FlpO mouse with a transgenic Flp-dependent reporter mouse. CRHR1 and reporter molecules were identified using immunocytochemistry and visualized via confocal microscopy in several brain regions in which CRHR1 expression and function is established.

**Results:** Expression of Flp-dependent viral constructs was highly specific to CRHR1-expressing cells in all regions examined (over 90% co-localization). In accord, robust and specific expression of the Flp-dependent transgenic reporter was observed in a reporter mouse, recapitulating endogenous CRHR1 expression.

**Conclusions:** The *Crhr1*-FlpO mouse enables selective genetic access to CRHR1-expressing cells within the mouse brain. When combined with Cre-lox or site-specific recombinases, the mouse facilitates intersectional manipulations of CRHR1-expressing neurons.

## Introduction

Corticotropin-releasing hormone (CRH) is an evolutionarily conserved peptide that functions as a critical molecular regulator of neuroendocrine, autonomic, and behavioral stress responses (Vale et al., 1981; Brown et al., 1985; Smith & Vale, 2006; Korosi & Baram, 2008; Joëls & Baram, 2009; Deussing & Chen, 2018). CRH plays a critical role in modulating the hypothalamic-pituitary-adrenal (HPA) axis, increasing adrenocorticotropic hormone (ACTH) and glucocorticoid levels, and stimulating autonomic nervous system activity (Vale et al., 1981; Smith & Vale, 2006; Bale & Vale, 2004). Within the brain, CRH is released from axonal terminals (Chen et al., 2001; Chen et al., 2004) and engages with its two G protein-coupled receptors, CRH receptor type 1 (CRHR1; Chang et al., 1993; Chen et al., 1993; Vita et al., 1993) and type 2 (CRHR2; Kishimoto et al., 1995; Lovenberg et al., 1995; Perrin et al., 1995), acting as a neuromodulator influencing neuronal and circuit function (Gunn et al., 2017; Baumgartner, Schulkin, & Berridge, 2021; George et al., 2012; Lemos, Shin & Alvarez, 2019; Garcia et al., 2014). The majority of known CRH functions within the brain are mediated via activation of CRHR1 (Chen et al., 2006; Howerton et al., 2014; Refojo et al., 2011; Sztainberg et al., 2011; Müller et al., 2003; Wang et al., 2011; Chen et al., 2016; Dedic et al., 2018), influencing diverse behavioral and cognitive processes (Cui et al., 2013; Gilpin, Yu, & Kash, 2021; Hupalo et al., 2019; Kimbrough et al., 2017; Simpson et al., 2020; Rajamanickam & Justice, 2022).

CRHR1 is abundantly expressed throughout the brain in distinct regionally specific patterns (Chen et al., 2000). In the neocortex, for example, CRHR1 neurons are present throughout layers II-VI where they receive local input from GABAergic CRH interneurons (Yan et al., 1998; Van Pett et al., 2000; Chen et al., 2000; Kubota et al., 2011). Cortical CRHR1 regulates anxiety-like behaviors (Magalhaes et al., 2010) and mediates stress-induced cognitive dysfunction (Hupalo et al., 2019; Uribe-Mariño et al., 2016). In the hippocampus, CRHR1 is highly expressed by pyramidal cells in areas CA1 and CA3, where the receptor is located on cell bodies and dendrites (Chen et al., 2004; Chen et al., 2010; Andres et al., 2013; Cursano et al., 2021) and GABAergic interneurons provide local CRH input (Chen et al., 2004; Gunn et al., 2017; Gunn et al., 2019). Stress-induced CRH release in the hippocampus modulates learning and memory processes and can destroy dendritic spines (Gunn et al., 2017; Chen et al., 2010; Ivy et al., 2010). The receptor is also expressed in numerous subcortical regions, including the nucleus accumbens, where CRH input may be both local or arrive from long-range projections (Lemos et al., 2012; Walsh et al., 2014; Baumgartner et al., 2021; Pan et al., 2024 Itoga et al., 2019; Birnie et al., 2023). CRH-CRHR1 signaling in the accumbens has been shown to regulate reward and aversion behaviors in a context- and sex-dependent manner (Birnie et al., 2023; Lemos et al., 2012). For detailed mapping of CRHR1 anatomical distribution throughout the mouse brain, please see Chen et al., 2000, Van Pett et al., 2000, Justice et al., 2008, and the Allen Mouse Brain Atlas (https://mouse.brain-map.org/; Lein et al., 2007).

Here, we describe the generation of a novel transgenic mouse, the *Crhr1*-FlpO mouse, which utilizes the codon-optimized flippase (FlpO) recombinase system driven by the *Crhr1* promoter. We test the specificity of FlpO expression within CRHR1-expressing cells using two separate complementary methods. First, we inject Flp-dependent adeno-associated viruses (AAVs) into target brain regions and employ fluorescent immunocytochemistry (ICC) to examine whether Flp-dependent expression is predominantly restricted to CRHR1-expressing cells. Second, we cross the *Crhr1*-FlpO mouse with a transgenic Flp reporter mouse to determine if the expression of this Flp-driven transgenic reporter molecule (mCherry) corresponds to the anatomical distribution of CRHR1, via direct comparison with immunocytochemistry of endogenous CRHR1 using an established specific antiserum. Both methods demonstrate high specificity of the *Crhr1*-FlpO mouse driven expression of viral and transgenic constructs with high specificity in CRHR1-expressing cells.

## Materials and Methods

### I. Experimental Animals

All experimental procedures were approved by the University of California-Irvine Institutional Animal Care and Use Committee and were in accordance with National Institute of Health (NIH) guidelines. Animals were group housed in a temperature and humidity-controlled room in standard housing with a 12-hour light/dark cycle (lights on 7:00 a.m., lights off 7:00 p.m.). Food and water were provided *ad libitum*. Adult male and female *Crhr1*-FlpO mice (2-3 months of age) were used for all stereotaxic virus injection surgeries (n = 12; 4 per target brain region).

#### Generation of the Crhr1-FlpO mouse

The *Crhr1*-FlpO mouse was generated using a bacterial artificial chromosome (BAC) transgenesis approach similar to the generation of the *Crfr1-cre* mouse (Jiang, Rajamanickam, & Justice, 2018) and the CRF_1_:Cre rat (Weera et al., 2022). Briefly, E. Coli containing a BAC (rp24-239f10) that covers the entire *Crhr1* genomic locus (*Fig. 1A*) was transformed with a construct encoding FlpO-pGHpa-WPRE with 5’ and 3’ targeting sequences (*Fig. 1B*). Colonies were screened for accurate insertion by PCR. Targeting was designed to insert the gene encoding FlpO to replace the ATG at the *Crhr1* translation start site (*Fig. 1C*). Following recombination, we isolated a single BAC clone and used PCR and restriction enzyme laddering to confirm the integrity of the BAC. BAC DNA was purified, linearized, and injected into single cell oocytes that were then implanted into a CD-1(ICR) foster mother. Six BAC transgenic founders were identified by genotyping offspring using primers specific to the BAC gene.

**Fig. 1.**
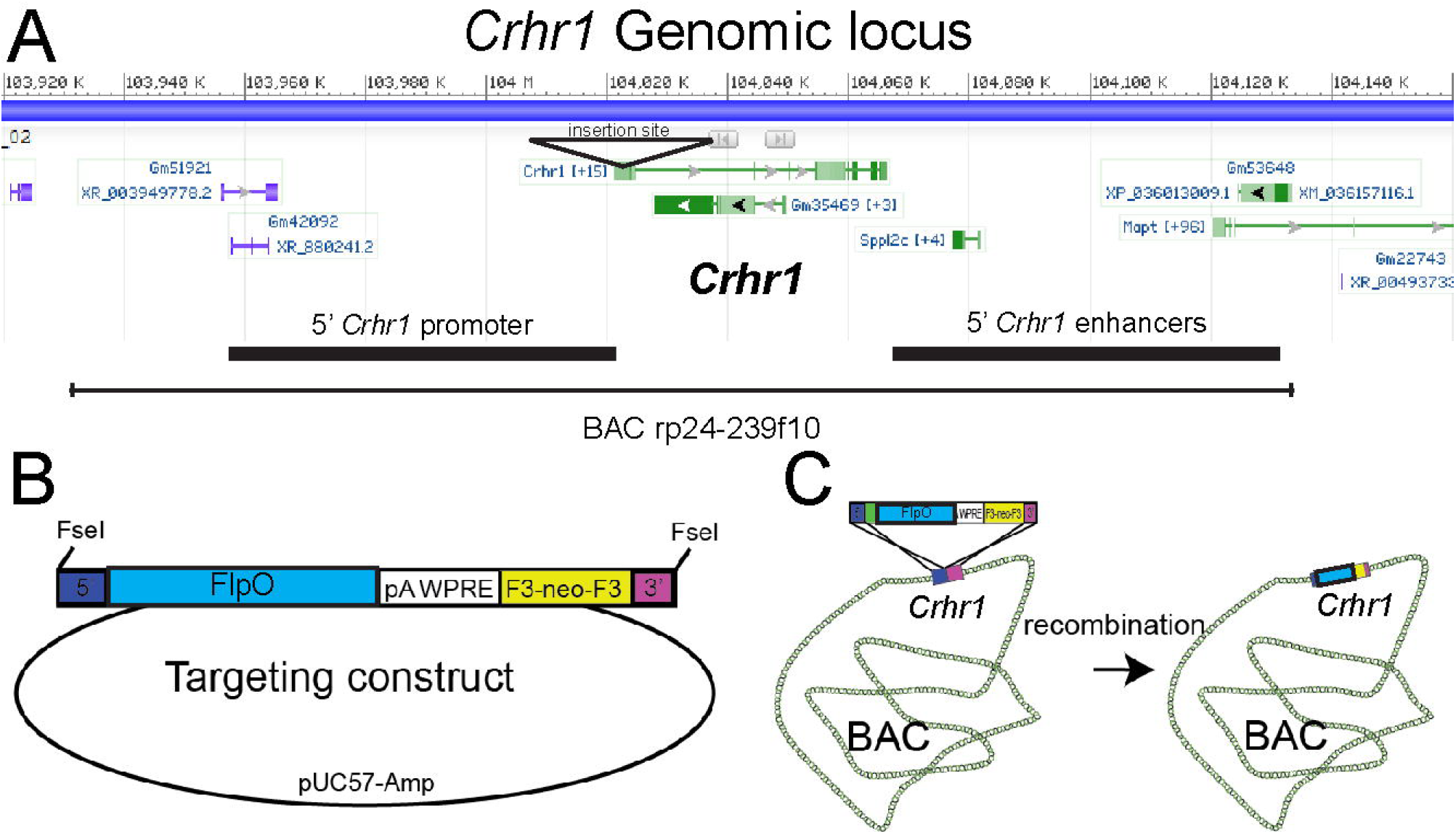
The *Crhr1*-FlpO BAC transgene construct. **(A)** *Crhr1* is located on chromosome 11 of the mouse. **(B)** The transgene, containing 5’ (blue) and 3’ (magenta) targeting sequences, FlpO recombinase, a 3’ polyA WPRE stabilizing sequence, and a F3-flanked neomycin resistance sequence (yellow). **(C)** Depicts the recombination process whereby the transgene was transformed into *E. coli* containing the rp24-239f10 BAC construct to integrate the transgene into the BAC, generating the product containing FlpO immediately upstream of the *Crhr1* translation ATG start site

Founders were backcrossed to C57/B6 mice, then outcrossed to a Rosa knock-in reporter mouse of Flp expression, RR1 (also referred to as FPi, kindly obtained from R. Ray; Gt(ROSA)25Sor^tm#(CAG-mCherry-hM4D)Rray^; JAX stock #029040; Ray et al, 2011, Lusk et al, 2022). This reporter allele is an intersectional genetics allele that expresses DREADD-Gi in the presence of both Cre and Flp recombinases, and mCherry in the presence of only Flp recombinase (shown in *Figs. 5-6*). The pattern of mCherry expression was compared with reported expression patterns of CRHR1 (Chen et al, 2000; Van Pett et al., 2000, Justice et al., 2008). Two lines displayed expression patterns very similar to CRHR1, and to each other. One of these was expanded and used for further analysis (*Crhr1*-FlpO).

### II. Viruses and Surgical Procedures

#### Viral vectors

Adeno-associated virus (AAV) vectors, AAV8-nEF-C_off_/F_on_-ChR2(ET/TC)-EYFP (Addgene viral prep # 137141-AAV8) and AAV8-nEF-C_off_/F_on_-ChR2-mCherry (Addgene viral prep # 137144-AAV8), were purchased from Addgene. These viral vectors were a gift to Addgene from Karl Deisseroth & INTRSECT 2.0 Project (Fenno et al., 2020). The viral titer of each virus was 2.6 × 10^13^ genotypic copies per milliliter (GC/mL).

#### Stereotaxic virus injections

Prior to the start of surgery, mice (PND 60-90) were anesthetized in an isoflurane chamber (4% isoflurane with 1.0 L/min oxygen flow) using a tabletop isoflurane vaporizer system. Mice were then placed on a robotic stereotaxic frame (Neurostar) with their head secured and maintained under anesthesia with a constant flow of 1.5% isoflurane (1.0 L/min oxygen flow rate). The analgesic Buprenorphine (0.1 mg/kg) was administered subcutaneously, and the shaved scalp was sterilized with iodine and 70% ethanol. An incision was made in the scalp and the skull was positioned flat along the anteroposterior and mediolateral axes, using Bregma and Lambda as landmarks. A dental drill was used to drill through the skull above target injection regions. Flp-dependent viral reporters expressing either ChR2-EYFP or ChR2-mCherry were loaded into a pulled glass pipette which was lowered into the brain at target coordinates. Viruses were injected in a volume of ∼200 nl per hemisphere at a rate of ∼20-40 nl per minute using a Picospritzer III apparatus set to 5-10 ms pulses. To prevent backflow of virus, the glass pipette was left in the injection site for 10 minutes after infusion. Post-surgery, mice were removed from anesthesia and placed in a clean cage on a heat pad and monitored until normal movement was observed (∼15-30 minutes). Coordinates for each of the targets were as follows: Dorsal hippocampus (AP -1.94, ML +/-1.40, DV 1.40) cortex (prefrontal/motor; AP +1.4, ML +/-0.5, DV 1.2), and nucleus accumbens (AP +1.2, ML +/-0.7, DV 4.5). Following viral injection into target regions, the virus was left to express for 4-6 weeks prior to perfusion and tissue processing.

### III. Perfusion and Sectioning

Mice were deeply anesthetized and euthanized with a lethal Euthasol (1% pentobarbital sodium and phenytoin sodium) then perfused with saline (0.9% NaCl in double-distilled H2O) followed by 4% paraformaldehyde (PFA) dissolved in 0.1 M phosphate buffer (PB, pH 7.4). Brains were removed and post-fixed in the same 4% PFA fixative overnight, followed by sequential overnight incubation steps in 15% sucrose and 25% sucrose (prepared with 0.1M PB, pH 7.4), all at +4°C. The brains were dried and frozen on dry ice for 10 minutes, then stored at -80°C. Brains were sectioned using a Leica CM1900 cryostat (Leica Microsystems, Germany). Prior to sectioning, tissue was placed in the cryostat set to -20°C for at least 1 hour to equilibrate to cryostat temperature. Brains were then placed on a cryostat mounting disk and encased in Tissue-Tek

O.C.T. Compound (Sakura, Ref #4583). Coronal sections, 25 μm thick, were acquired in a series of six (∼150 μm between one section to the next in each series). Serial sections were stored at -20°C in an antifreeze solution.

### III. Immunocytochemistry (ICC)

#### Antibody Characterization

Goat anti-CRHR1 (Everest Biotech, CAT# EB08035; RRID:AB_2260976) is a polyclonal antibody that recognizes the N-terminus of CRHR1 (amino acids 107-117). This N-terminus sequence in CRHR1 resides in the first extracellular domain and is distinct from the CRHR2 peptide sequence (Wille et al., 1999; Perrin et al., 2006). Western blots of human, mouse, and rat tissue lysates produce immunoblot bands of the appropriate size (manufacturer’s datasheet; Baglietto-Vargas et al., 2015), matching the molecular weight we previously reported with a separate antibody for CRHR1 targeting the C-terminus (Chen et al., 2000). ICC on free-floating tissue using this antibody strongly recapitulates the cellular characteristics and brain-wide distribution patterns of CRHR1 (Chen et al., 2000; van Pett et al., 2000) and transgenic CRHR1 reporter mice (Justice et al., 2008).

Chicken anti-GFP (Aves Labs, CAT# GFP-1020; RRID:AB_10000240) is a polyclonal antibody that recognizes green fluorescent protein (GFP; derived from jellyfish *Aequorea victoria*) and its variants, including YFP (manufacturer’s datasheet). Western blots using this antibody recognize GFP and GFP fusion proteins (Zhang et al., 2023). This antibody has been successfully used on free-floating brain slices to label viral-mediated GFP in mouse neurons (Oyola et al., 2017).

Rabbit anti-RFP (Rockland, CAT# 600-401-379; RRID:AB_2209751) is a polyclonal antibody that recognizes the entire amino acid sequence of red fluorescent protein (RFP; derived from mushroom anemone *Discoma*) and its variants, including mCherry and tdTomato (manufacturer’s datasheet). This antibody recognizes both mCherry and tdTomato on free-floating brain sections (Kooiker et al., 2023; Yu et al., 2018; Gunn et al., 2019).

#### ICC on free-floating brain tissue sections

We performed sequential fluorescent immunolabeling of target antigens on free-floating brain tissue sections to visualize CRHR1 and Flp-dependent reporter molecules (EYFP or mCherry) in single cells (see Table 1). Briefly, sections were washed and permeabilized in 0.01 M PBS containing 0.3% Triton X-100 (PBS-T) for 30 minutes (3 × 10 minutes). Sections were then treated with 0.3% H_2_O_2_ in PBS-T for 30 minutes followed by washing with PBS-T (2 × 15 minutes). To block non-specific binding of secondary antibodies, sections were incubated in 5% normal serum derived from the same host species as secondary antibodies (rabbit or donkey; NRS or NDS) for 1 hour. Next, sections were incubated at +4°C for 72 hours in primary antibody Goat anti-CRHR1 (1:2,000). Secondary antibody labeling was then performed at room temperature, in the dark, with agitation using either Cy3 AffiniPure Rabbit Anti-Goat (1:500; Jackson Laboratories, CAT# 305-165-003) or Donkey anti-Goat Alexa Fluor 488 (Invitrogen, CAT# A-11055) to label CRHR1 with green-or red-conjugated fluorophores, respectively. After washing in PBS-T (3 × 5 minutes), sections were incubated in either primary antibody Chicken anti-GFP (1:5000; targeting EYFP) or Rabbit anti-RFP (1:2000; targeting mCherry) at +4°C for 72 hours. This was followed by washing and application of secondary antibody Alexa Fluor 488 AffiniPure Rabbit Anti-Chicken (1:500; Jackson Laboratories, CAT# 303-545-003) or Donkey anti-Rabbit Alexa Fluor 568 (1:400; Invitrogen; CAT#A-10042) for 2 hours at room temperature in the dark with agitation. All antibodies were diluted in PBS-T (0.3% Triton X-100). After wet-mounting on microscope slides, sections were cover-slipped with mounting medium for fluorescent staining (Bioenno Lifesciences, CAT# 032019).

**Table 1.**
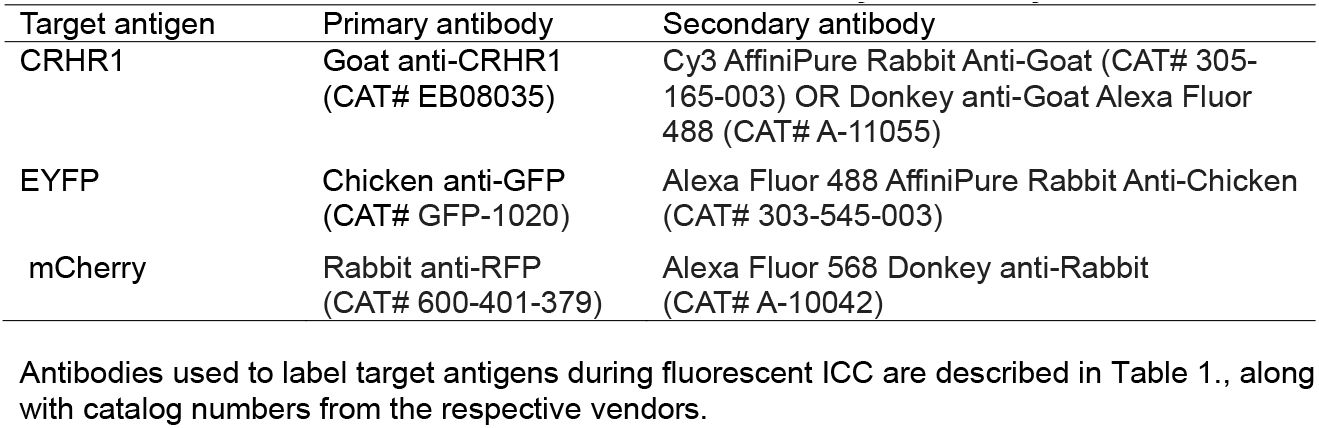
List of antibodies used for fluorescent immunocytochemistry.

An additional set of sections was stained for CRHR1 only using standard avidin-biotin-complex (ABC) methods. Sections were washed and treated with 0.3% H_2_O_2_ as described above. These sections were then blocked with 2% NRS for 30 minutes, rinsed in PBS-T for 10 minutes, and incubated in Goat anti-CRHR1 (1:5000) for 48 hours at +4°C. Following primary antibody incubation, sections were incubated in biotinylated rabbit anti-goat IgG (1:200, Vector, Burlingame, CA) for 1 hour at room temperature with agitation. After washing (3 × 5 minutes) in PBS-T, these sections were incubated in ABC solution (1:100) at room temperature for 2 hours followed by washing (3 × 5 minutes). These sections were then treated with 0.04% 3,3-diaminobenzidine (DAB) with 0.5% nickel chloride for 8-10 minutes.

### IV. Confocal Microscopy

Tissues with fluorescently labeled antigens were imaged using a Zeiss LSM-510 Confocal Microscope. Alexa Fluor 488-conjugated secondary antibodies (green emission spectrum) were visualized using an Argon 488 nm excitation laser and a 500-530 nm emission filter. Cyanine3 (Cy3)- and Alexa Fluor 568-conjugated secondary antibodies (red emission spectrum) were visualized using a He/Ne 543 nm excitation laser and a 560-615 nm emission filter. Z stack images were acquired at intervals of 1 μm to determine the presence or absence of colocalization through single cells. Brightness, contrast, and sharpness of images were adjusted in ImageJ to aid visualization.

### V. Data Analysis

We employed unbiased sampling of sections to examine the colocalization of endogenous CRHR1 and Flp-dependent viral reporters (EYFP or mCherry), using one in six serial sections as above. Because viral injection is not expected to infect all CRHR1 cells, we limited the analyses to specificity and did not attempt to assess sensitivity. Therefore, individual cells were categorized as either colocalizing (containing signal for both the virus and CRHR1; ‘specific’) or non-colocalizing (containing signal for the virus only; ‘non-specific’). The percentage of cells within the examined brain regions that co-expressed both CRHR1 and Flp-dependent viral reporter molecules was calculated by the fraction of colocalizing cells over the total number of virus-labeled cells.

## Results

### The *Crhr1-*FlpO mouse

The generation of the *Crhr1*-FlpO mouse is described in the Materials and Methods and depicted in *Figure 1*. Viable, fertile mice with stable BAC transmission were generated and employed throughout the current studies.

### Colocalization of Flp-dependent viral reporters with endogenous CRHR1

#### Cortex

Flp-dependent viral reporters were injected into the cortex of *Crhr1*-FlpO mice (n = 4; 2 male and 2 female). *Figure 2* shows representative data from a *Crhr1*-FlpO mouse injected with AAV8-nEF-C_off_/F_on_-ChR2-mCherry (*Fig. 2A*). Endogenous CRHR1 (shown in green) is distributed in a laminar pattern through cortical layers II-VI (*Fig. 2B & 2C)*. Colocalization analyses of endogenous CRHR1 and the viral reporters (shown in red) indicated a robust overlap (*Fig. 2B-2E*). Viral expression was concentrated primarily in pyramidal cells within layers II-III, IV, and V, in accordance with the expression pattern of endogenous CRHR1 (*Fig. 5; also see* Chen et al., 2000; Van Pett et al., 2000). In the cortex, viral reporters were occasionally present in layer VI and rarely found in layer I. Quantifying cells co-expressing both CRHR1 and viral reporters vs. the total cells expressing viral reporters revealed high specificity of Flp-mediated expression in CRHR1-immunoreactive cells: 90.4% and 95.3% of virus-expressing cells also expressed CRHR1 in males and females, respectively.

**Fig. 2.**
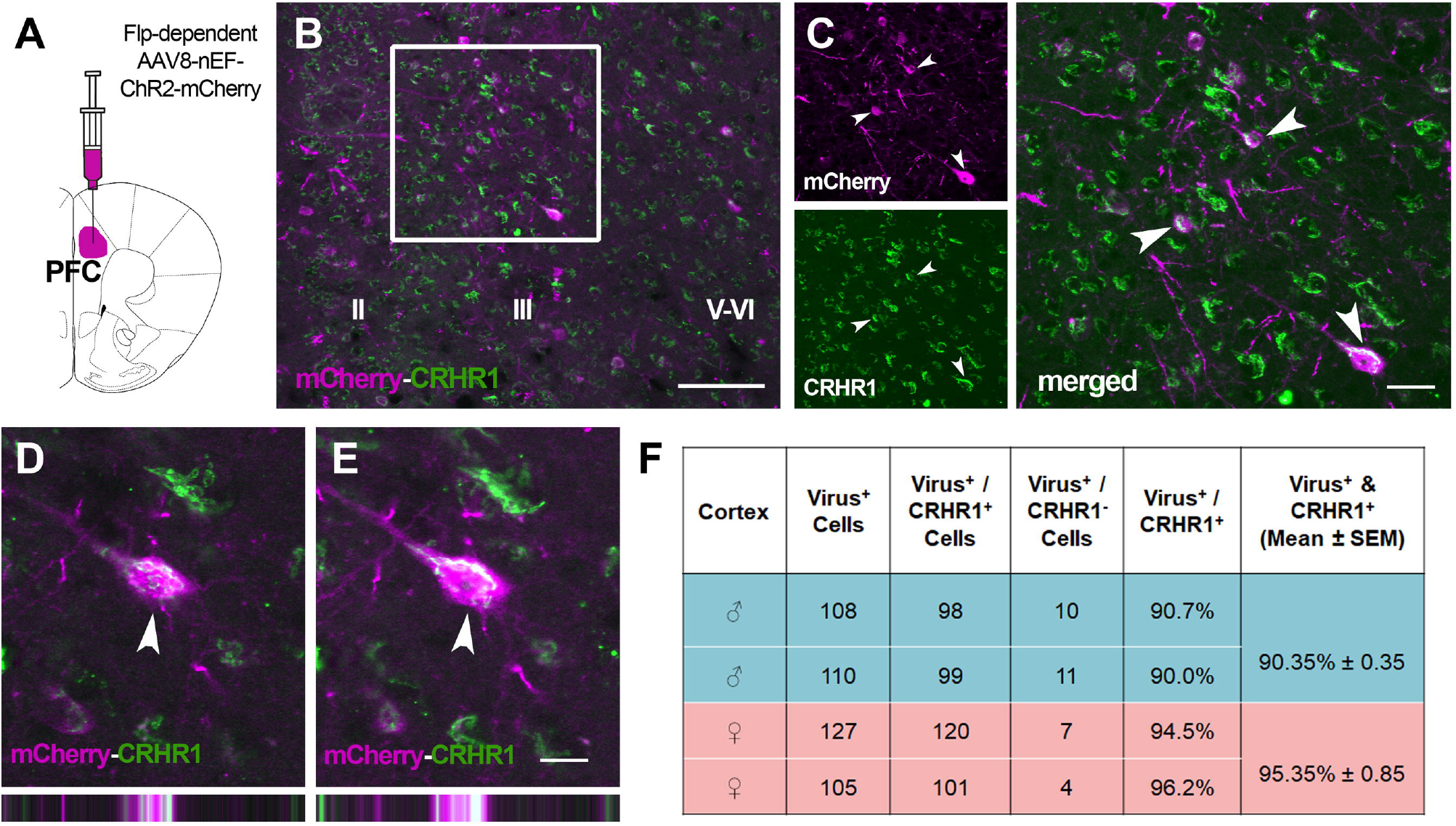
Flp-dependent virus-derived reporter molecules colocalize with endogenous CRHR1 in the prefrontal cortex. **(A)** Schematic depicting PFC virus injection. **(B)** Image of the cortex showing endogenous CRHR1 expression (green) and Flp-dependent mCherry (magenta). Scale bar = 100 µm. **(C)** Higher magnification of the boxed region in panel A shows mCherry (top) and CRHR1 (bottom) channels separate and merged (right). Scale bar = 25 µm. **(D & E)** Higher magnification showing labeling of mCherry and CRHR1 within a single neuron at 1 µm intervals in the Z plane, demonstrating colocalization of both target antigens. Bars on the bottom show orthogonal views of the same neuron. Scale bar = 12.5 µm **(F)** Quantification of the percentage of virus-labeled cells in the cortex that also co-express CRHR1 vs. cells which express virus only in individual male and female mice

#### Dorsal hippocampus

We injected Flp-dependent viruses into the dorsal hippocampus of *Crhr1*-FlpO mice, targeting the CA1 region (n = 4; 2 male and 2 female). *Figure 3* shows representative data from a mouse injected with AAV8-nEF-C_off_/F_on_-ChR2-EYFP in the dorsal hippocampus (*Fig. 3A*). Virus-labeled cells (shown in green) were identified in CA1 (*Fig. 3B & 3C*), CA3, the dorsal subiculum and, less frequently, the stratum oriens and stratum radiatum. Immunolabeling of virus-driven fluorophores and CRHR1 (shown in red) strongly overlapped (*Fig. 3D & 3E)*, with 97.1% of reporter-expressing cells co-expressing CRHR1 in females and 92.2% in males (*Fig. 3F*).

**Fig. 3.**
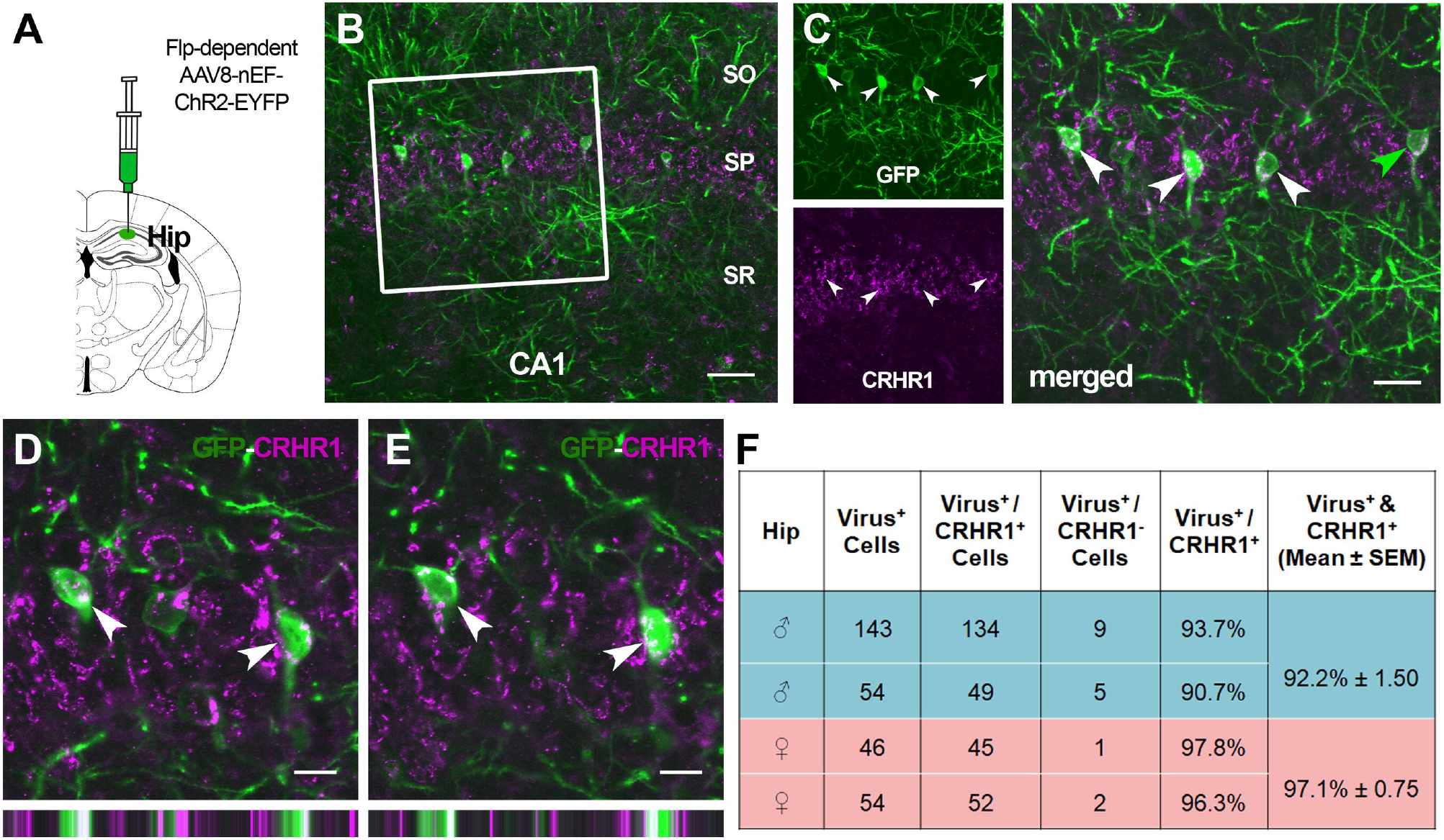
Flp-dependent virus-derived reporter molecules colocalize with endogenous CRHR1 in the dorsal hippocampus. **(A)** Schematic depicting dorsal hippocampus virus injection. **(B)** Image of the dorsal hippocampus CA1 region showing endogenous CRHR1 expression (magenta) and Flp-dependent EYFP (green). Scale bar = 50 µm. **(C)** Higher magnification of the boxed region in panel A shows EYFP (top) and CRHR1 (bottom) channels separate and merged (right). Scale bar = 25 µm **(D & E)** Higher magnification showing labeling of EYFP and CRHR1, demonstrating colocalization of both target antigens. Images show two different Z planes of the same neurons. Bars on the bottom show orthogonal views of the same neurons. Scale bars = 12.5 µm. **(F)** Quantification of the percentage of virus-labeled cells in the dorsal hippocampus that also co-express CRHR1 vs. cells which express virus only. Abbreviations: SO, stratum oriens; SP, stratum pyramidale; SR, stratum radiatum

#### Nucleus accumbens

We injected Flp-dependent viruses into the nucleus accumbens of *Crhr1*-FlpO mice (*Fig. 4A*; n = 4; 2 male and 2 female). *Figure 4* shows representative data from a mouse injected with AAV8-nEF-C_off_/F_on_-ChR2-EYFP. Virus-labeled cells (shown in green) were identified in the accumbens medial shell (*Fig. 4B*) and core. The vast majority of viral Flp reporter-expressing cells co-expressed endogenous CRHR1 (shown in red; *Fig. 4C & 4D*). Specifically, 95.4% and 94.3% of virus-expressing cells co-expressed CRHR1 in males and females, respectively (*Fig.4E*).

**Fig. 4.**
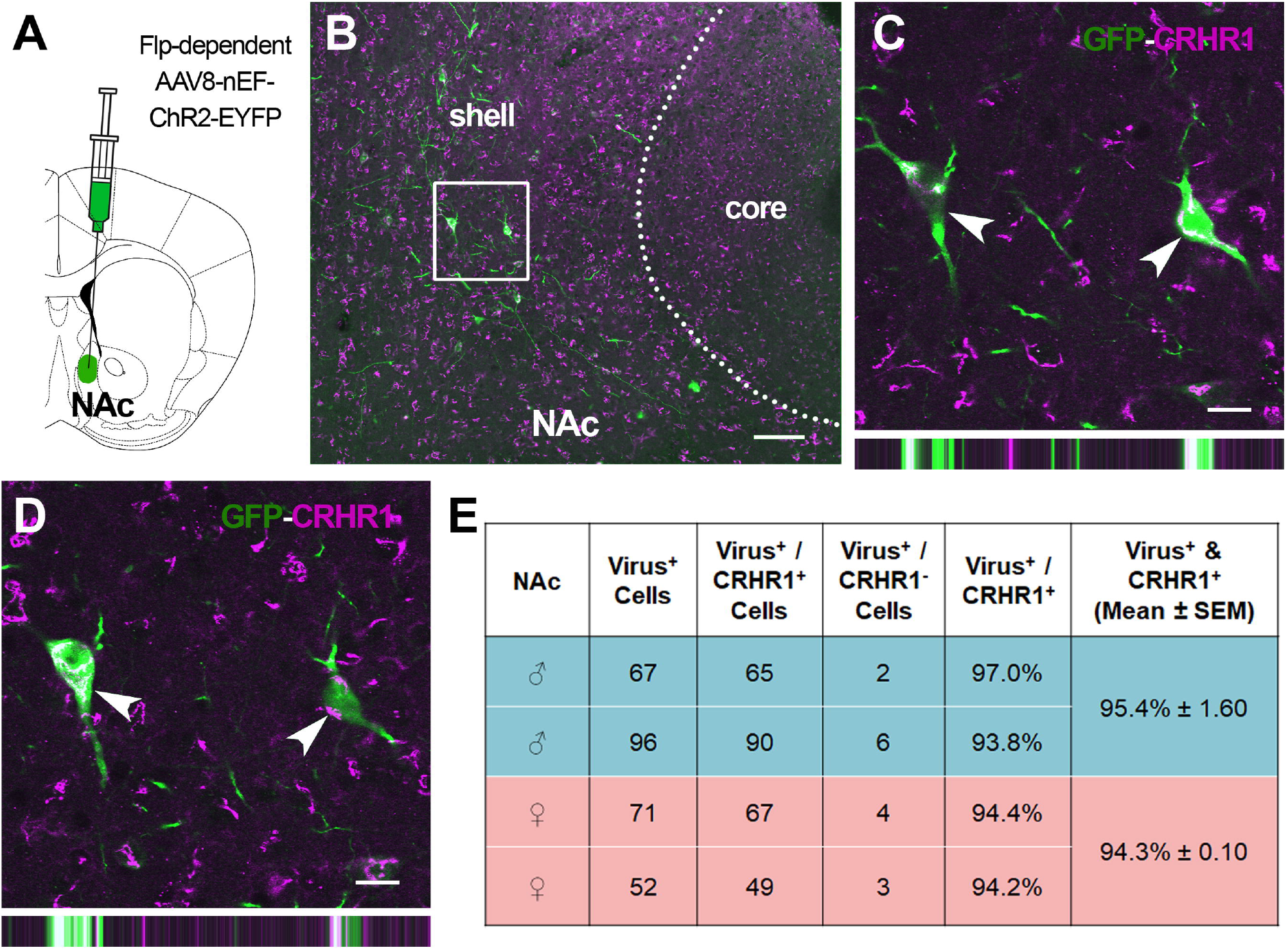
Flp-dependent virus-derived reporter molecules colocalize with endogenous CRHR1 in the nucleus accumbens. **(A)** Schematic showing nucleus accumbens (NAc) virus injection. **(B)** Image of the nucleus accumbens showing endogenous CRHR1 expression (magenta) and Flp-dependent EYFP (green). Scale bar = 100 µm. **(C & D)** Higher magnification images of the boxed region in panel B shows colocalization of EYFP and CRHR1 within single neurons at 1 µm intervals in the Z plane. Images show two different Z planes of the same neurons. Bars on the bottom show orthogonal views of the same neurons. Scale bar = 15 µm. **(E)** Quantification of the percentage of virus-labeled cells in the nucleus accumbens that also co-express CRHR1 vs. cells which express virus only in individual male and female mice

### Flp reporter expression in *Crhr1*-FlpO x RR1 mice recapitulates anatomical patterns of endogenous CRHR1 distribution

To increase the rigor of our analyses, we used an independent method that does not rely upon viral transfection. We crossed the *Crhr1*-FlpO mouse with the Flp-driven transgenic reporter mouse RR1, enabling visualization of the distribution of FlpO recombinase throughout the whole mouse brain via the constitutive expression of Flp-dependent mCherry in the resulting offspring. We then compared the anatomical distribution of mCherry to that of endogenous CRHR1 determined using immunocytochemistry.

In the cortex, mCherry was abundantly distributed throughout cortical layers II-VI (*Fig 5A &* 5C), in accord with expression patterns of endogenous CRHR1 (*Fig. 5B & 5D*). Layers IV/V, followed by layers II/III, had the highest density of mCherry-expressing cells (Table 2). Similarly, the distribution of mCherry throughout the anatomical subdivisions of the amygdala strikingly overlapped that of endogenous CRHR1 (*Fig. 6A & 6B*). Thus, the basolateral amygdala (BLA) was defined by numerous large and densely packed cells. The lateral amygdala (LA) also contained numerous cell clusters, whereas the central nucleus medial portion (CeAm) was characterized by a moderate number of lower density cells. The central nucleus lateral division (CeAl) and the basomedial amygdala (BMA) both contained sparser, widely separated cells relative to the rest of the amygdala. In these subdivisions, mCherry-positive cells were fewer than cells expressing endogenous CRHR1. This might be attributable to either small differences in the specific anteroposterior location of the sampled sections or limited *Crhr1* promoter-driven FlpO expression in these neurons. Lastly, we assessed the molecular layer of the dentate gyrus as a negative control region (*Supplemental Fig. 1*), where we could not detect CRHR1-immuno-reactivity or Flp-dependent mCherry signal.

**Table 2.** Density of cells expressing the RR1 reporter in a *Crhr1*-FlpO x RR1 mouse.

| Region | RR1 reporter density | CRHR1 neuron density |
| --- | --- | --- |
| Neocortex |  |  |
| Layer I | – | – |
| Layer II/III | ++ | ++/+++ |
| Layer IV/V | +++ | +++ |
| Layer VI | + | +++ |
| Retrosplenial cortex |  |  |
| Layer I | – | – |
| Layer II/III | ++ | ++ |
| Layer V | + | ++/+++ |
| Amygdala |  |  |
| Lateral amygdala | +++ | +++ |
| Basolateral amygdala | +++ | +++ |
| Central nucleus, lateral portion | + | +++ |
| Central nucleus medial portion | ++ | ++ |
| Basomedial amygdala | + | +++ |
| Molecular layer of the dentate gyrus | – | – |

Density scores are based on the number of reporter (mCherry) or CRHR1 positive cells and the proportion of positive cells that are contiguous with other positive cells. – no positive cells; +, a few cells; ++, many cells, mixture of contiguous and non-contiguous cells; +++, many positive cells, most or all cells are contiguous.

**Fig. 5.**
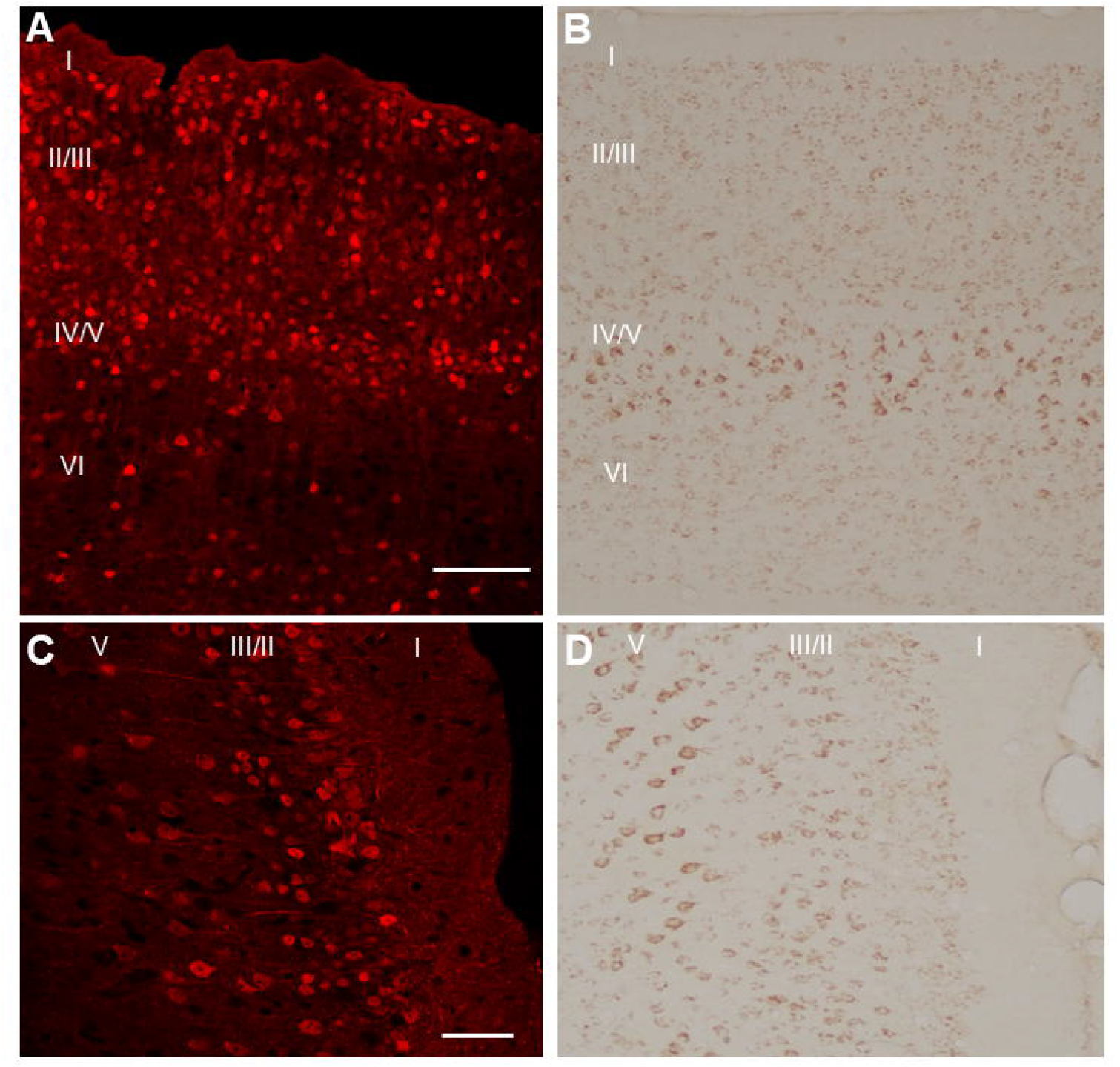
Flp-dependent reporter expression in *Crhr1*-FlpO x RR1 mice recapitulates endogenous CRHR1-immunoreactivity patterns in the cortex. **(A)** Flp-dependent mCherry expression (red) in the neocortex of *Crhr1*-FlpO x RR1 mice. Scale bar = 100 µm. **(B)** Corresponding image shows endogenous CRHR1 (brown) immunoreactivity in the neocortex. **(C)** Flp-dependent mCherry expression in the retrosplenial cortex of *Crhr1*-FlpO x RR1 mice. Scale bar = 50 µm. **(D)** Corresponding image of endogenous CRHR1 in the retrosplenial cortex

**Fig. 6.**
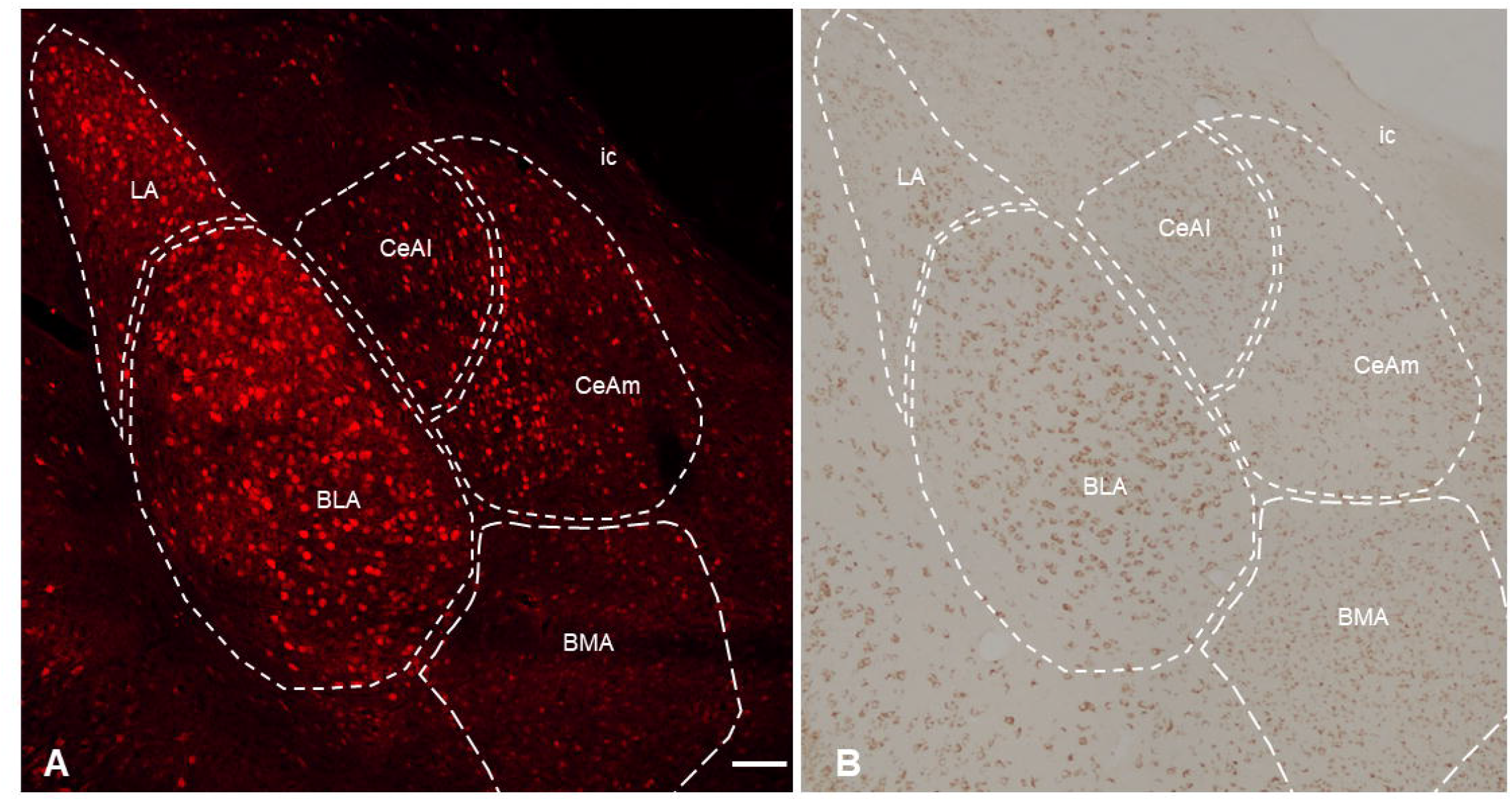
Flp-dependent reporter expression in *Crhr1*-FlpO x RR1 mice recapitulates endogenous CRHR1-immunoreactivity patterns in the amygdala. **(A)** Flp-dependent mCherry expression (red) in the amygdala of *Crhr1*-FlpO x RR1 mice. Scale bar = 100 µm. **(B)** Image shows endogenous CRHR1 (brown) immunoreactivity throughout the amygdala. Abbreviations: LA, Lateral amygdala; BLA, basolateral amygdala; CeAl, central nucleus lateral portion; CeAm, central nucleus medial portion; BMA; basomedial amygdala.

## Discussion

We developed and validated a new transgenic mouse, *Crhr1*-FlpO, that allows for genetic access and manipulation of CRHR1-expressing cells. The experiments presented here, focusing on three salient CRHR1-expressing brain regions (i.e., the cortex, hippocampus, and nucleus accumbens), demonstrate that this mouse allows for Flp-dependent expression of genetic constructs that is highly specific to CRHR1-expressing cells

In the cortex, Flp-dependent virus reporter molecules and CRHR1-expressing neurons were distributed in a laminar pattern throughout layers II-VI. As with the distribution of endogenous CRHR1 in the cortex, Flp reporters were most densely located in layers II/III and IV/V. In the dorsal hippocampus, reporter molecules and CRHR1-expressing neurons were localized predominantly to pyramidal neurons in layers CA1, CA3, and the dorsal subiculum. The stratum oriens and stratum radiatum also contained sparse virus and CRHR1 labeling. Lastly, in the nucleus accumbens, Flp-dependent reporter molecules and CRHR1-expressing neurons were present throughout both the accumbens medial shell and core. In all three brain regions, Flp-dependent reporter molecules were found to express with high specificity in cells that also expressed CRHR1. Limitations of the study are implicit in the limited range and quantity of infection by the virus: only a minority of endogenous CRHR1 expressing neurons co-expressed the viral reporters. Thus, we report on the specificity but not the sensitivity of the mouse.

Crossing the *Crhr1*-FlpO mouse with the RR1 Flp-dependent reporter line enabled a comprehensive assessment of Flp expression in this mouse in a manner that is independent of the transfection efficacy and diffusion of injected viruses. Offspring resulting from this cross express mCherry in a pattern that is highly consistent with endogenous CRHR1 expression. We also crossed *Crhr1*-FlpO mice with another available Flp reporter line, Ai65F. This cross resulted in significantly fewer reporter (tdTomato) positive cells (*Supplemental Fig. 2)*. This finding is consistent with Zhao and colleagues’ (2023) observations of poor reporter expression when crossing the *CRH-FlpO* mouse with Ai65F compared to other Flp-dependent reporter lines. The low expression levels may be a result of differences in the design of the FRT flanked stop cassette. Another explanation is relatively low expression levels of FlpO driven by the *Crhr1* promoter. Indeed, the level of promoter activity driving FlpO expression as a limiting factor in the efficacy of Flp-mediated recombination has been well-characterized (Zhao et al., 2023). Zhao and colleagues demonstrate that acute restraint stress improves FlpO-mediated recombination in the *CRH-FlpO* mouse, attributed to elevated activity of the *Crh* promoter driving FlpO expression. Dynamic expression of CRHR1 is thus an important consideration, as stress modulates the activity of the *Crhr1* promoter and correspondingly the production of FlpO in this mouse (Van Pett et al., 2000; Greetfeld et al., 2009; Meng et al., 2011). Additionally, cells labeled in our reporter cross will comprise not only those that express CRHR1 in adulthood but also any cell that transiently expressed CRHR1 during development.

These limitations, coupled with the variability in the degree of cell labeling using viral injection, prevent quantitative assessments of the *sensitivity* of the *Crhr1*-FlpO transgenic mouse.

Notably, to manipulate CRHR1-activity, transfection of a large proportion of neurons will likely be required, involving optimization of viral titer, expression time, and potentially different AAV serotypes. Indeed, this mouse may not be ideal for broad quantitative mapping of the CRHR1 expression pattern, and other CRHR1 transgenic reporter mice may be more suited for this use (Justice et al 2008; Jiang et al., 2018). Rather, the *Crhr1*-FlpO mouse is highly *specific* and is thus an excellent tool used in combination with Cre lines where selective expression of virally delivered transgenes needs to be specified to two distinct cell types. In addition, combining *Crhr1*-FlpO with other Cre lines can be used to specify expression in cells that express both CRHR1 and another gene, when using Cre- and Flp-dependent viral constructs.

Thus, while Cre-driver lines have been developed for targeting CRHR1-expressing cells (Dedic et al., 2018; Jiang, Rajamanickam, & Justice, 2018), a key advantage of the *Crhr1*-FlpO mouse is that it is compatible in genetic crosses with mice that rely upon Cre-lox systems. The advent of transgenic mouse models led to the widespread implementation of Cre recombinase across a diversity of transgenic lines (Birnie et al., 2023; Kooiker et al., 2023; Taniguchi et al., 2011; Navabpour, Kwapis, & Jarome, 2020; Cui et al., 2021). Conversely, Flp-driver lines were less quickly adopted, mainly because the original derivatives of Flp recombinase had weaker transduction efficiency compared to the Cre-lox system (Ringrose et al., 1998). However, FlpO, a codon-optimized form of FlpE that is more thermostable in the mammalian body, achieves transduction rates more comparable to Cre recombinase (Raymond & Soriano, 2007; Kranze et al., 2010). This makes novel FlpO transgenic lines powerful tools that complement the already robust Cre-dependent transgenic systems. The crossing of Flp- and Cre-driver lines will give researchers more refined control over neurons and circuits, facilitating access to and manipulation of multiple cellular populations within the same animal.

In sum, the *Crhr1*-FlpO mouse is a reliable tool to target and manipulate CRHR1-expressing cells in the brain. Research on CRH-CRHR1 signaling has often capitalized on the direct application of CRH or CRH receptor blockers, cell culture, electrophysiology, immunocytochemistry, and models of stress (Gunn et al., 2017; Curran et al., 2017; Blank et al., 2002; Schierloh et al., 2007;Lemos et al., 2012). The implementation of Cre-driver lines to support visualization (Justice et al., 2008; Chen et al., 2015; Vandael et al., 2021; Walsh et al., 2014) and genetic manipulation (Birnie et al., 2023; Jiang et al., 2018; Kratzer et al., 2013; Flandreau et al., 2015; Hooper et al., 2018) of CRH or CRHR1-expressing neuronal populations has greatly enhanced our understanding of the functional consequences of CRH-mediated signaling throughout the brain. We propose that the *Crhr1*-FlpO mouse will be a valuable addition to this arsenal of genetic tools, particularly when used in a synergistic fashion with existing transgenic mouse lines (e.g., CRH-IRES-Cre; Taniguchi et al., 2011) and tools for sensing calcium and neurotransmitter dynamics *in* vivo (Nakai, Ohkura, & Imoto, 2001; Sun et al., 2020).

## Conclusion

We describe and validate a novel *Crhr1*-FlpO mouse as a reliable specific tool that provides genetic access to CRHR1-expressing cell populations. Along with extensive preclinical research demonstrating the importance of CRHR1 signaling across behavioral and physiological domains, the CRH-CRHR1 system is critically implicated in mental health and disease (Bale et al., 2010; Binder & Nemeroff, 2010; Hupalo et al. 2019; Stanton, Price & Manning, 2023; Demarchi, Pawluski, & Bosch, 2021; Weera & Gilpin, 2019; Russell & Lightman, 2019; Mantsch et al., 2016; Spanagel, Noori, & Heilig, 2014). The synergistic utilization of genetic tools for dissecting CRHR1-mediated signaling and circuit activity in will enhance our investigations into the mechanisms that underlie the function and disruption of endogenous stress systems, thereby advancing our knowledge of healthy and disordered brain states.

## Supporting information

Flp-dependent reporter expression in Crhr1-FlpO x RR1 mice demonstrates lack of endogenous CRHR1-immunoreactivity in the dentate molecular layer

Flp-dependent tdTomato expression in Crhr1-Flp x Ai65F mice

## Acknowledgments

The authors thank Dr. M. Birnie and Dr. A. Floriou-Servou for insightful comments. The work was supported by the National Institutes of Health, RO1 MH132680 (T.Z.B), R56 MH114032 (N.J.), R01 MH112768 (N.J.), T32 DA50558 (M.H.), the Bren Foundation and the Welch Foundation.

## Statements & Declarations

### Author Contributions

MH, NJ, and TZB conceptualized and designed the study. Figure preparation, data collection and analysis were performed by Mason Hardy and Yuncai Chen. The first draft of the manuscript was written by Mason Hardy. Writing, reviewing, and editing was performed by Mason Hardy, Tallie Z. Baram, and Nicholas J. Justice. All authors read and approved the final manuscript.

### Data Availability

The data used to support the findings of this study are available from the corresponding author upon reasonable request.

### Ethics approval

All housing and experimental procedures were performed in accordance with the National Institute of Health (NIH) guidelines and approved by the University of California-Irvine Institutional Animal Care and Use Committee.

