## Supplementary figures and images for "Targeting Corticotropin-Releasing Hormone Receptor Type 1 (CRHR1) Neurons: Validating the Specificity of a Novel Transgenic *Crhr1*-FlpO Mouse"

### Flp-dependent reporter expression in Crhr1-FlpO x RR1 mice demonstrates lack of endogenous CRHR1-immunoreactivity in the dentate molecular layer

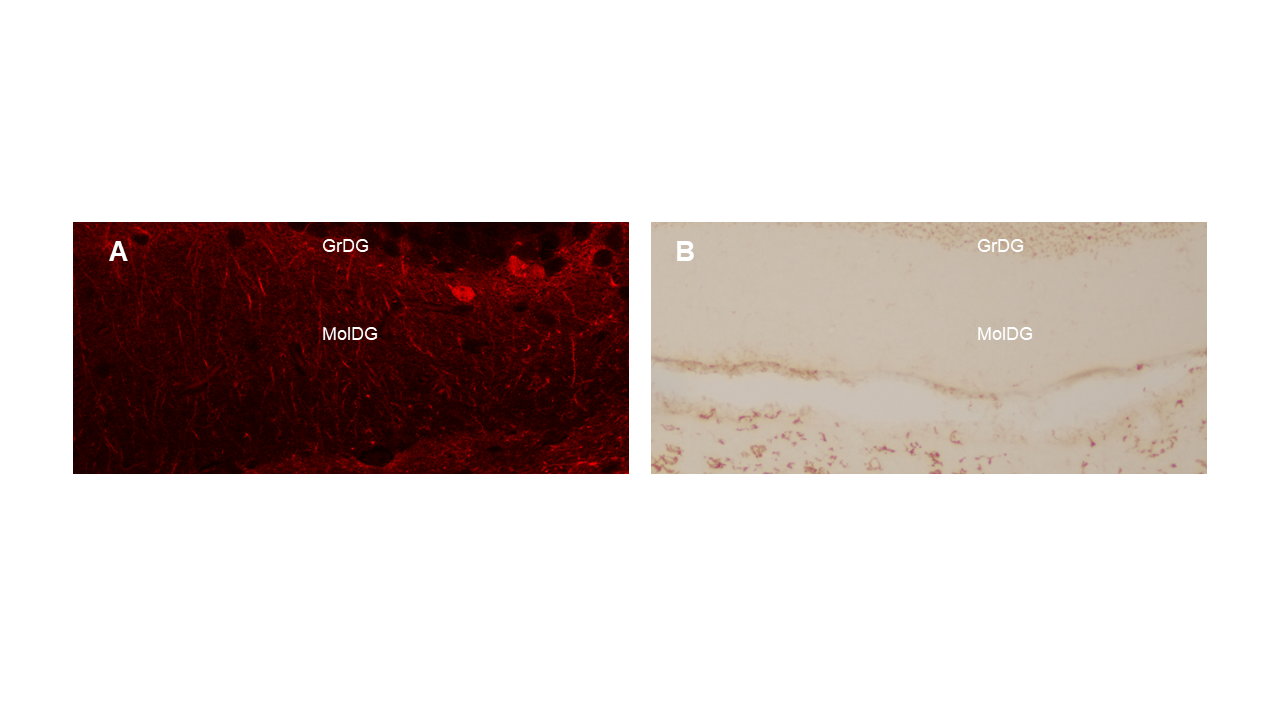

### Flp-dependent tdTomato expression in Crhr1-Flp x Ai65F mice

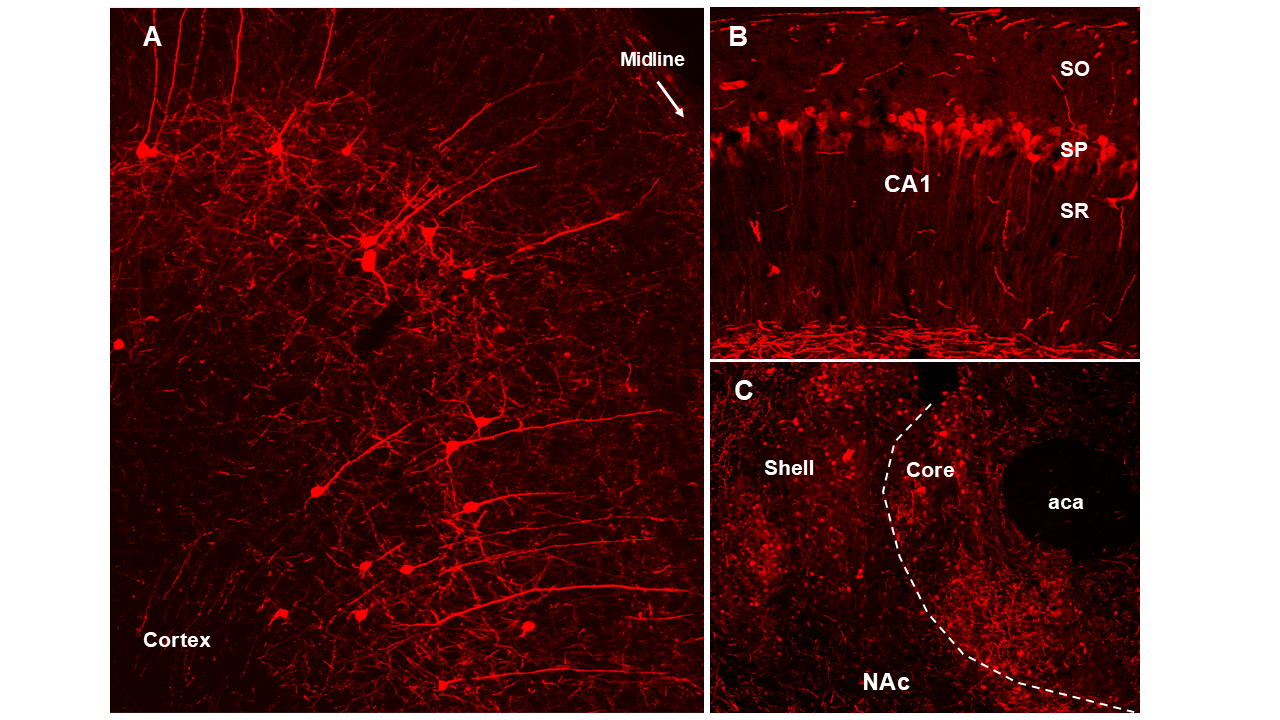
